# Dynamic changes in the brain protein interaction network correlates with progression of Aβ42 pathology in *Drosophila*

**DOI:** 10.1101/501213

**Authors:** Harry M. Scholes, Adam Cryar, Fiona Kerr, David Sutherland, Lee A. Gethings, Johannes P. C. Vissers, Jonathan G. Lees, Christine A. Orengo, Linda Partridge, Konstantinos Thalassinos

## Abstract

Alzheimer’s disease (AD), the most prevalent form of dementia, is a progressive and devastating neurodegenerative condition for which there are no effective treatments. Understanding the molecular pathology of AD during disease progression may identify new ways to reduce neuronal damage. Here, we present a longitudinal study tracking dynamic proteomic alterations in the brains of an inducible *Drosophila melanogaster* model of AD expressing the Arctic mutant Aβ42 gene. We identified 3093 proteins from flies that were induced to express Aβ42 and age-matched healthy controls using label-free quantitative ion-mobility data independent analysis mass spectrometry. Of these, 228 proteins were significantly altered by Aβ42 accumulation and were enriched for AD-associated processes. Network analyses further revealed that these proteins have distinct hub and bottleneck properties in the brain protein interaction network, suggesting that several may have significant effects on brain function. Our unbiased analysis provides useful insights into the key processes governing the progression of amyloid toxicity and forms a basis for further functional analyses in model organisms and translation to mammalian systems.

## Introduction

Alzheimer’s disease (AD) is a progressive and devastating neurodegenerative disease that is the most prevalent form of dementia [1]. Symptoms initially present as episodic memory loss and subsequently develop into widespread cognitive impairment. Two brain lesions are pathological hallmarks of the disease: plaques and neurofibrillary tangles. Plaques are extracellular aggregates of amyloid beta (Aβ) [2], whereas, neurofibrillary tangles are intraneuronal aggregates of hyperphosphorylated tau [3,4]. In addition to these hallmarks, the AD brain experiences many other changes, including metabolic and oxidative dysregulation [5,6], DNA damage [7], cell cycle re-entry [8], axon loss [9] and, eventually, neuronal death [6,10].

Despite a substantial research effort, no cure for AD has been found. Effective treatments are desperately needed to cope with the projected increase in the number of new cases as a result of longer life expectancy and an ageing population. Sporadic onset is the most common form of AD (SAD), for which age is the major risk factor. Familial AD (FAD)—a less common (<1%), but more aggressive, form of the disease—has an early onset of pathology before the age of 65 [11]. FAD is caused by fully penetrant mutations in the Aβ precursor protein (APP) and two subunits—presenilin 1 and presenilin 2—of the Ɣ-secretase complex that processes APP in the amyloidogenic pathway to produce Aβ. Whilst the exact disease mechanisms of AD are not yet fully understood, this has provided support for Aβ accumulation as a key player in its cause and progression [1]. Aβ42—a 42 amino acid variant of the peptide—is neurotoxic [12], necessary for plaque deposition [13] and sufficient for tangle formation [14]. The Arctic mutation in Aβ42 (Glu22Gly) [15] causes a particularly aggressive form of familial AD that is associated with an increased rate and volume of plaque deposition [16]. Genetic analyses of SAD, however, suggest a complex molecular pathology, in which alterations in neuro-inflammation, cholesterol metabolism and synaptic recycling pathways may also be required for Aβ42 to initiate the toxic cascade of events leading to tau pathology and neuronal damage in dementia.

Comparison of proteomic analyses of post-mortem human brains have further revealed an increase in metabolic processes and reduction in synaptic function in AD [17]. Oxidised proteins also accumulate at early stages in AD brain, probably as a result of mitochondrial ROS production [18], and redox proteomic approaches suggest that enzymes involved in glucose metabolism are oxidised in mild cognitive impairment and AD [19,20]. Moreover, phospho-proteomic approaches have revealed alterations in phosphorylation of metabolic enzymes and kinases that regulate phosphorylation of chaperones such as HSP27 and crystallin alpha B [21]. Of note, however, there is little proteomic overlap between studies using post-mortem human brain tissue, which may reflect the low sample numbers available for such studies, differences in comorbidities between patients and confounding post-mortem procedures [17]. Although valuable, post-mortem studies also reflect the end-stage of disease and, therefore, do not facilitate measurement of dynamic alterations in proteins as AD progresses.

Animal models of AD, generated through transgenic over-expression of human APP or tau, provide an opportunity to track proteomic alterations at pre- and post-pathological stages, thus facilitating insight into the molecular mechanisms underlying disease development and revealing new targets for drugs to prevent AD progression. Analyses of transgenic mice models of AD have revealed some overlapping alterations in metabolic enzymes, kinases and chaperones with human AD brain [17]. Only one study, however, has tracked alterations in protein carbonylation over time, showing increases in oxidation of metabolic enzymes (alpha-enolase, ATP synthase α-chain and pyruvate dehydrogenase E1) and regulatory molecules (14-3-3 and Pin1) in correlation with disease progression [22].

Adult-onset *Drosophila* models of AD have been generated by over-expressing human Aβ42 peptide exclusively in adult fly neurons using inducible expression systems. These models have been shown to develop progressive neurodegenerative phenotypes, such as reduced climbing ability, and shortened lifespan [23]. Taking advantage of the short lifespan of the fly, and the flexible nature of the inducible model, we have performed a longitudinal study of the brain proteome to capture the effects of Aβ42-toxicity in the brain from the point of induction and across life. We identified 3093 proteins using label-free quantitative ion-mobility data independent analysis mass spectrometry (IM-DIA-MS) [24], 1854 of which were common to healthy and Aβ42 flies. Of these, we identified 228 proteins that were significantly altered in AD, some of which overlapped with normal ageing but the majority of which were ageing-independent. Proteins altered in response to Aβ42 were enriched for AD processes and have statistically significant network properties in the brain protein interaction network. We also show that these proteins are likely to be bottlenecks for signalling in the network, suggesting that they comprise important proteins for normal brain function. Our data is a valuable resource to begin to understand the dynamic properties of Aβ42 proteo-toxicity during AD progression. Future functional studies will be required to determine the causal role of these proteins in mediating progression of AD using model organisms and to translate these findings to mammalian systems.

## Results

### Proteome analysis of healthy and Aβ42-expressing fly brains

Using an inducible transgenic fly line expressing human Arctic mutant Aβ42 (TgAD) [23] (Fig 1A), we confirmed a previously observed [23] reduction in lifespan following Aβ42 induction prior to proteomic analyses (Fig 1B).

**Figure 1.**
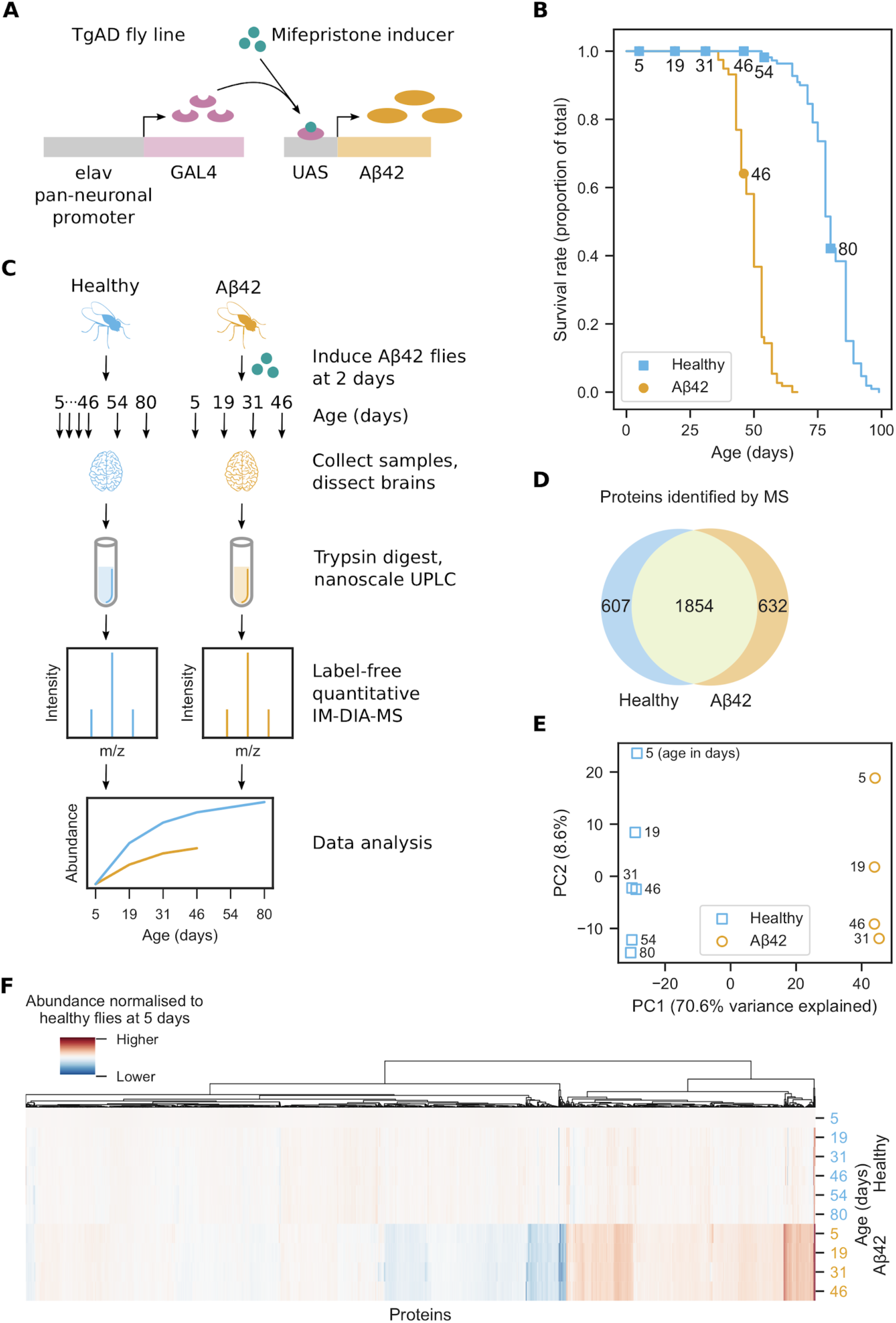
Proteome analysis of healthy and AD fly brains. (**A**) *Drosophila melanogaster* transgenic model of AD (TgAD) that expresses Arctic mutant Aβ42 in a mifepristone-inducible GAL4/UAS expression system under the pan-neuronal elav promoter. (**B**) Survival curves for healthy and Aβ42 flies. Aβ42 flies were induced to express Aβ42 at 2 days. Markers indicate days that MS samples were collected. (**C**) Experimental design of the brain proteome analysis. Aβ42 flies were induced to express Aβ42 at 2 days. For each of the three biological repeats, 10 healthy and 10 Aβ42 flies were collected at 5, 19, 31 and 46 days, as well as 54 and 80 days for healthy flies. Proteins were extracted from dissected brains and digested with trypsin. The resulting peptides were separated by nanoscale liquid chromatography and analysed by label-free quantitative IM-DIA-MS. (**D**) Proteins identified by IM-DIA-MS. (**E**) Principal component analysis of the IM-DIA-MS data. Axes are annotated with the percentage of variance explained by each principal component. (**F**) Hierarchical biclustering using relative protein abundances normalised to their abundance in healthy flies at 5 days.

To understand how the brain proteome is affected as Aβ42 toxicity progresses, fly brains were dissected from healthy and Aβ42 flies at 5, 19, 31 and 46 days, and at 54 and 80 days for healthy controls, then analysed by label-free quantitative IM-DIA-MS (Fig 1C, Supplementary Data 1). 1854 proteins were identified in both healthy and Aβ42 fly brain from a total of 3093 proteins (Fig 1D), which is typical for recent fly proteomics studies [25,26].

For the 1854 proteins identified in both healthy and Aβ42 flies, we assessed the reliability of our data. Proteins were highly correlated between technical and biological repeats (Fig S1). We used principal component analysis of the protein abundances to identify sources of variance (Fig 1E). Healthy and Aβ42 samples are clearly separated in the first principal component, probably due to the effects of Aβ42. In the second principal component, samples are separated by increasing age, due to age-dependent or disease progression changes in the proteome. These results show that whilst ageing does contribute to changes in the brain proteome (8.7% of the total variance), much larger changes are due to expression of Aβ42 (70.6%) and this may reflect either a correlation with the ageing process or progression of AD pathology. We confirmed this result using hierarchical biclustering of protein abundances in Aβ42 versus healthy flies at 5 days (Fig 1F). The results reveal that most proteins do not vary significantly in abundance with age in healthy flies, but many proteins are differentially abundant in Aβ42 flies.

### Analysis of brain proteome dysregulation in Aβ42 flies

We next identified the proteins that were significantly altered following Aβ42 expression in the fly brain. To achieve this, we used five methods commonly used to analyse time course RNA-Seq data [27] and classified proteins as significantly altered if at least two methods detected them [28]. We identified 228 significantly altered proteins from 740 proteins that were detected by one or more methods (Fig 2A). A comparison of popular RNA-Seq analysis tools [29] showed that edgeR [30] has a high false positive rate and variable performance on different data sets, whereas, DESeq2 [31] and limma [32] have low false positive rates and perform more consistently. We observed a similar trend in our data set. limma and DESeq2 detected the lowest number of proteins, with 21 proteins in common (Fig S2A). edgeR detected more proteins, of which 38 were also detected by DESeq2 and 16 by limma. EDGE [33] and maSigPro [34] detected vastly more proteins, 464 of which were only detected by one method. Principal component analysis shows that edgeR, DESeq2 and limma detect similar proteins, whereas, EDGE and maSigPro detect very different proteins (Fig S2B).

**Figure 2.**
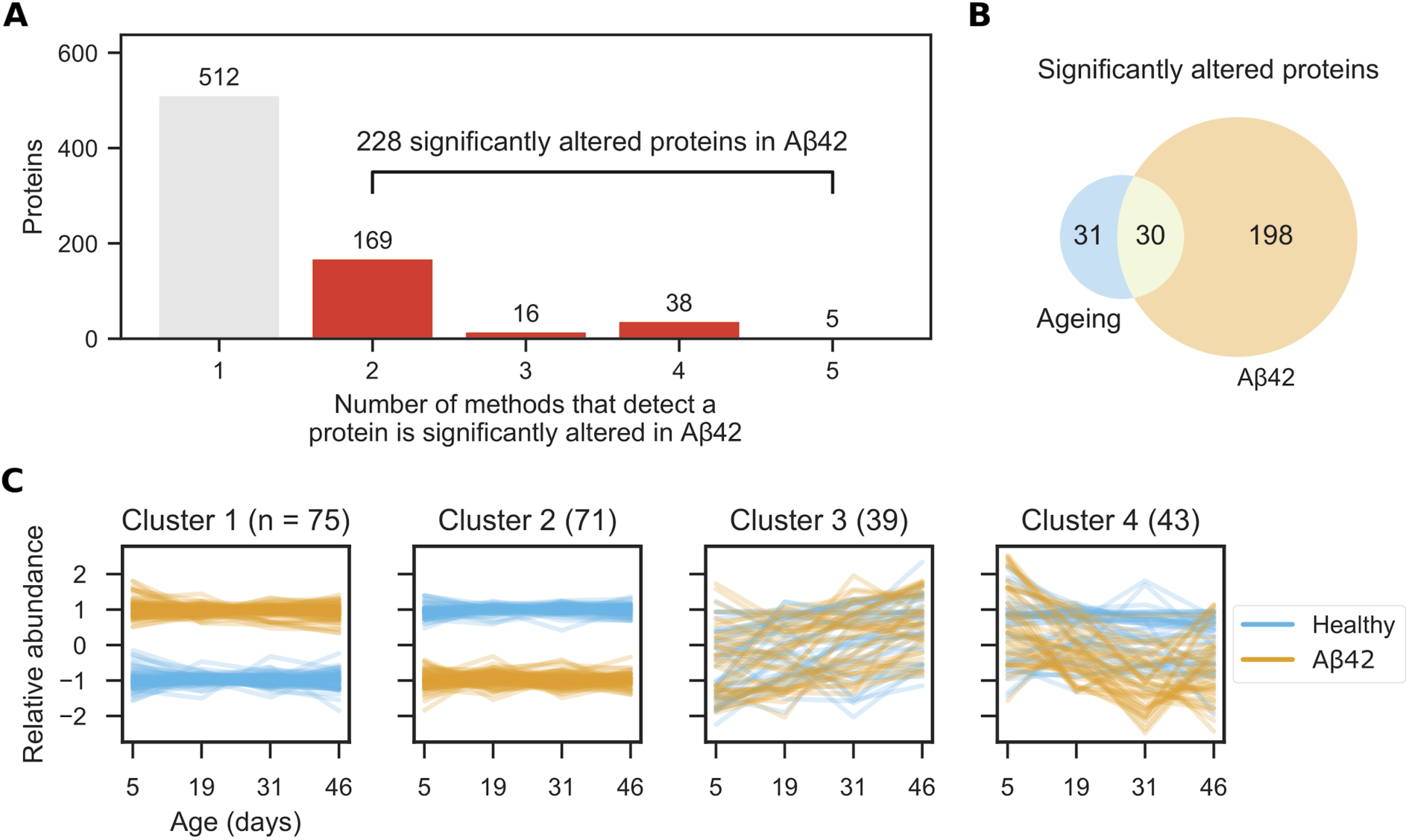
Brain proteome dysregulation in AD. (**A**) Proteins significantly altered in AD were identified using five methods (EDGE, edgeR, DESeq2, limma and maSigPro) and classified as significantly altered if at least two methods detected them. (**B**) Significantly altered proteins in AD (from **A**) and ageing. (**C**) Significantly altered protein abundances were *z* score-transformed and clustered using a Gaussian mixture model.

Although these methods should be able to differentiate between proteins that are altered in Aβ42 flies from those that change during normal ageing, we confirmed this by analysing healthy flies separately. In total, 61 proteins were identified as significantly altered with age (Fig S3), of which 30 were also identified as significantly altered in AD (Fig 2B) and 31 in normal ageing alone. These proteins are not significantly enriched for any pathways or functions. Based on our results, we concluded that the vast majority of proteins that are significantly altered in AD are not altered in normal ageing and that AD causes significant dysregulation of the brain proteome.

To understand the dynamics of protein alterations following Aβ42 induction, we clustered the profiles of proteins significantly altered in Aβ42 flies using a Gaussian mixture model (Fig 2C). The proteins clustered best into four sets (Fig S4). In comparison to healthy flies, cluster 1 contains proteins that have consistently higher abundance in Aβ42 flies. Conversely, cluster 2 contains proteins that have lower abundance in Aβ42 flies. The abundances of proteins from clusters 1 and 2 are affected from the onset of disease at day 5, and remain at similar levels as the disease progresses. Dysregulation of these proteins may initiate AD pathogenesis, be involved in early stages of disease progression, or represent defense mechanisms that could be harnessed for protection. Proteins in cluster 3 follow a similar trend in healthy and Aβ42 flies and increase in abundance with age. However, cluster 4 proteins decrease in abundance as the disease progresses, whilst remaining steady in healthy flies. Further work is required to determine whether reduction of these proteins plays a causal role in disease pathogenesis that could be targeted therapeutically, or whether their decline represents a protective response to damage.

We performed a statistical Gene Ontology enrichment analysis on each cluster, but found no enrichment of terms. Furthermore, we also saw no enrichment when we analysed all 228 proteins together.

### Brain proteins significantly altered by Aβ42 have distinct network properties

Following the analyses of brain proteome dysregulation in Aβ42 flies, we analysed the 228 significantly altered proteins in the context of the brain protein interaction network to determine whether their network properties are significantly different to the other brain proteins. Using a subgraph of the STRING [35] network induced on the 3093 proteins identified by IM-DIA-MS, we calculated four graph theoretic network properties (Fig 3A) of the 183 significantly altered proteins contained in this network: *degree*, the number of edges that a node has; *shortest path*, the smallest node set that connect any two nodes; *largest connected component*, the largest node set for which all nodes have at least one edge to any of the other nodes; and *betweenness centrality*, the proportion of all the shortest paths in the network that a particular node lies on.

**Figure 3:**
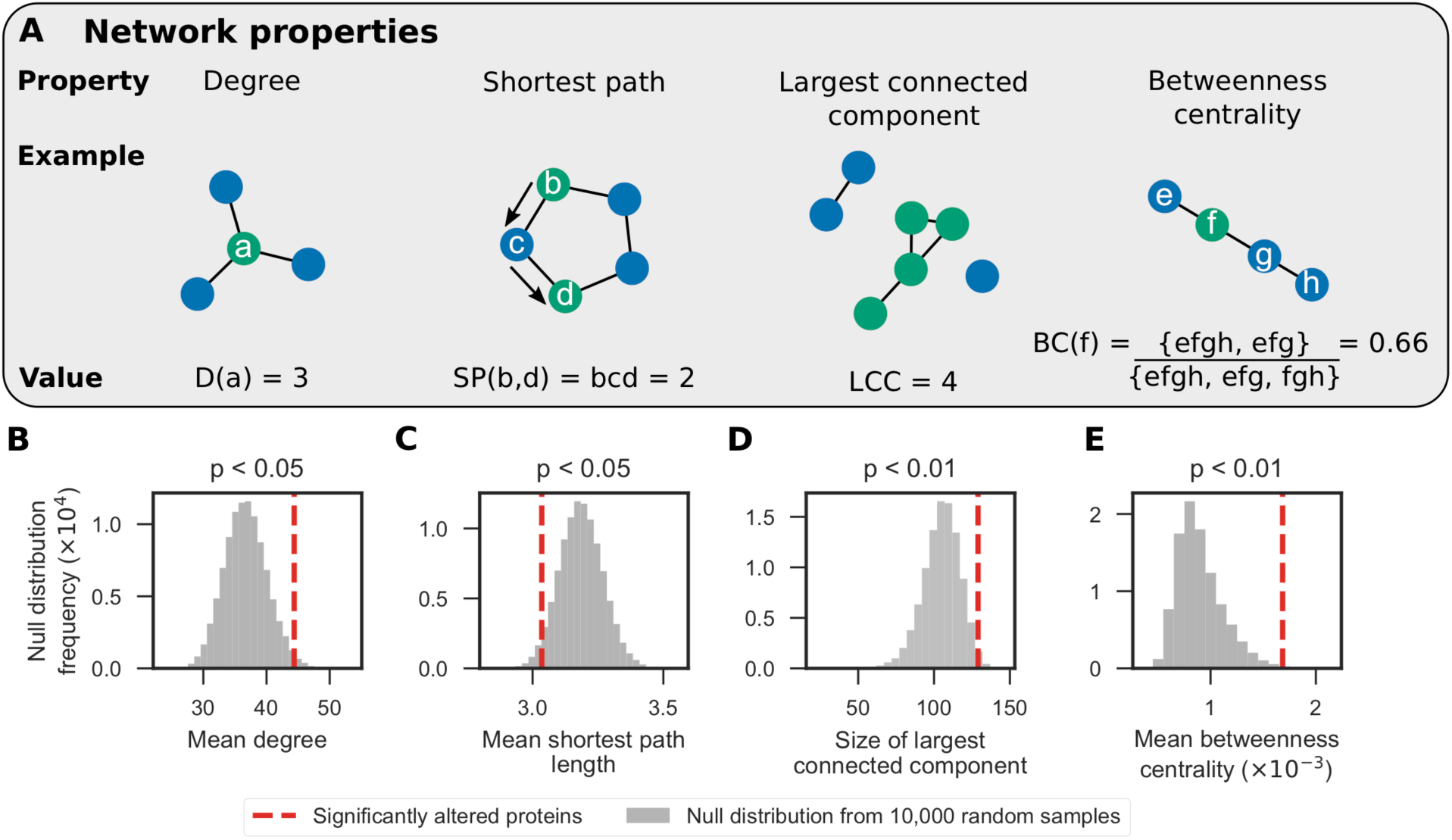
Significantly altered proteins have statistically significant network properties in the brain protein interaction network. **(A)** Network properties that were calculated: *degree*, the number of edges that a node has; *shortest path*, the smallest node set that connect any two nodes; *largest connected component*, the largest node set for which all nodes have at least one edge to any of the other nodes; and *betweenness centrality*, the proportion of all the shortest paths in the network that a particular node lies on. Using a subgraph of the STRING network induced on the 3093 proteins identified by IM-DIA-MS in healthy and Aβ42 flies, the significance of four network characteristics were calculated for the 183 significantly altered proteins contained in this subgraph. (B) mean degree; (C) mean shortest path length between a node and the remaining 182 nodes; (D) the size of the largest connected component in the subgraph induced on these nodes; and (E) mean betweenness centrality. Non-parametric p-values were calculated using null distributions of the test statistics, simulated by randomly sampling 183 nodes from the network 10,000 times.

We performed hypothesis tests and found that these proteins have statistically significant network properties. Firstly, the significantly altered proteins make more interactions than expected (mean degree p < 0.05; Fig 3B). Therefore, these proteins may further imbalance the proteome by disrupting the expression or activity of proteins they interact with. Secondly, not only are these proteins close to each other (mean shortest path p < 0.05; Fig 3C), but also 129 of them form a connected component (size of largest connected component p < 0.01; Fig 3D). These two pieces of evidence suggest that Aβ42 disrupts proteins at the centre of the proteome. Lastly, these proteins lie along shortest paths between many pairs of nodes (mean betweenness centrality p < 0.01; Fig 3E) and may control how signals are transmitted in cells. Proteins with high betweenness centrality are also more likely to be essential genes for viability [36]. Taken together, these findings suggest that the proteins significantly altered in AD are important in the protein interaction network, and that dysregulation of these proteins may have significant consequences for the brain proteome and therefore function.

### Predicting the severity of Aβ42-induced protein alterations using network properties

We predicted how severely particular Aβ42-associated protein alterations may affect the brain using two network properties—the tendency of a node to be a hub or a bottleneck. In networks, nodes with high degree are hubs for communication, whereas nodes with high betweenness centrality are bottlenecks that regulate how signals propagate through the network. Protein expression tends to be highly correlated to that of its neighbours in the protein interaction network. One exception to this rule, however, are bottleneck proteins, whose expression tends to be poorly correlated with that of its neighbours [36]. This suggests that the proteome is finely balanced and that the expression of bottleneck proteins is tightly regulated to maintain homeostasis. We analysed the hub and bottleneck properties of the significantly altered proteins and identified four hub-bottlenecks and five nonhub-bottlenecks that correlate with Aβ42 expression (Fig 4A) and analysed how their abundances change during normal ageing and as pathology progresses (Fig 4B).

**Figure 4.**
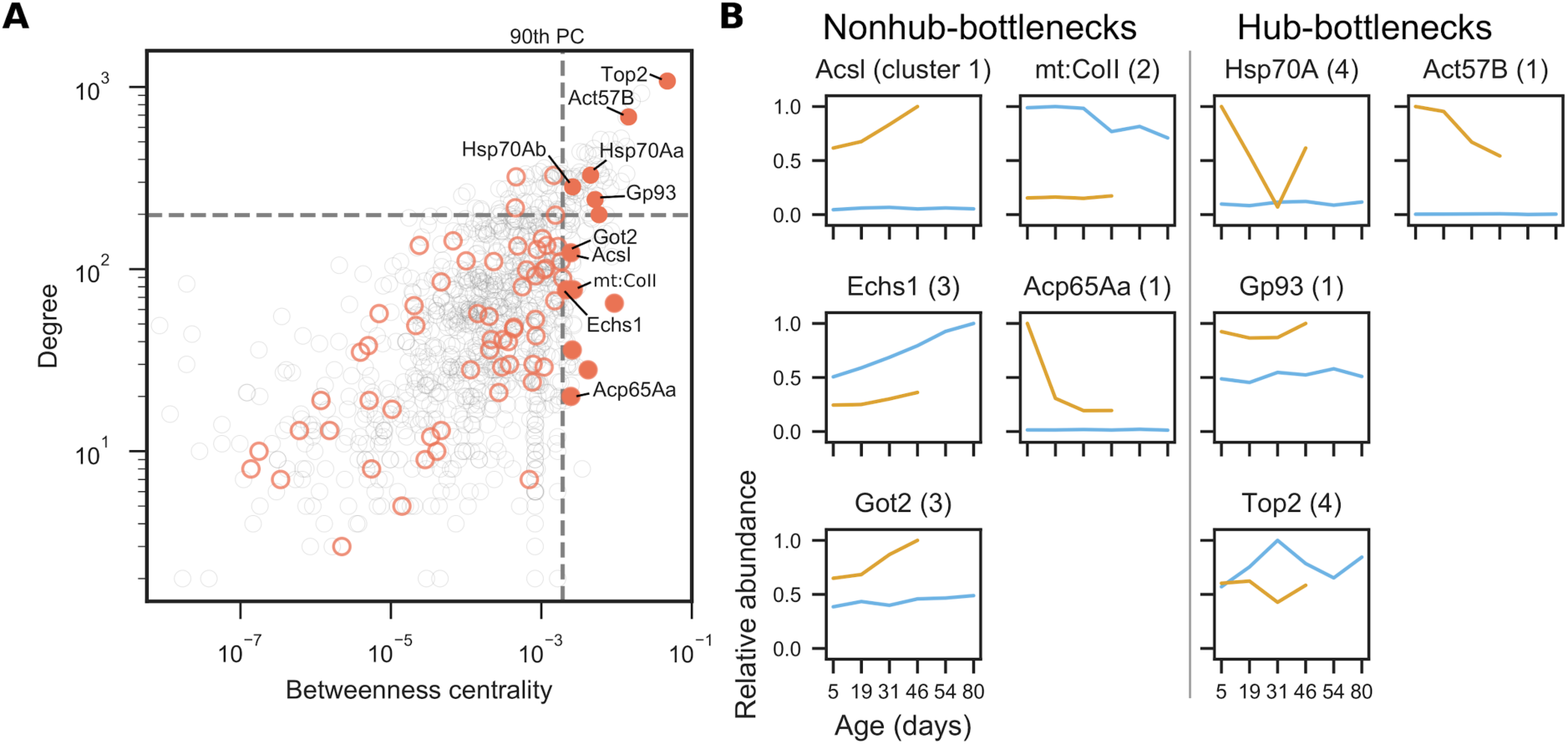
Analysis of hubs and bottlenecks in the brain protein interaction network. In networks, nodes with high degree are hubs and nodes with high betweenness centrality are bottlenecks. (**A**) Degree (hub-ness) is plotted against betweenness centrality (bottleneck-ness) in the brain protein interaction network for all proteins identified by IM-DIA-MS (grey circles). Of the significantly altered proteins (red circles), hub-bottleneck (> 90th percentile (PC) for degree and betweenness centrality) and nonhub-bottleneck proteins (> 90th PC for betweenness centrality) are highlighted (filled red circles). (**B**) Profiles of significantly altered bottleneck proteins implicated in Aβ42 toxicity. Maximum abundances are scaled to 1. Numbers in parentheses denote which cluster from Fig 2C the protein was in.

#### Nonhub-bottlenecks: Acs1, Echs1, Got2, mt:CoII and Acp65Aa

Three of the nonhub-bottlenecks, Acyl-CoA synthetase long chain (Acs1), Enoyl-CoA hydratase, short chain 1 (Echs1), and Aspartate aminotransferase (Got2), are metabolic enzymes with previous links to neuronal function and damage. Acs1 and Echs1 are involved in the production of acetyl-CoA from fatty acids. Many enzymes involved in acetyl-CoA metabolism associate with AD leading to acetyl-CoA deficits in the brain and loss of cholinergic neurons [6]. Got2 produces the neurotransmitter L-glutamate from aspartate, is involved in assembly of synapses and becomes elevated following brain injury [37]. Brain Acs1 and Got2 levels were stably expressed throughout normal ageing in our healthy flies but increased upon Aβ42 induction and continued to rise with age in Aβ42 flies. This suggests that levels of these proteins increase independently of ageing in AD but correlate closely with disease progression. On the other hand, Echs1 abundance increases in healthy flies during normal ageing, but its levels were reduced upon Aβ42 induction and its ageing-dependent increase was diminished in Aβ42 flies compared to controls. This may reflect a protective response with ageing that is suppressed by Aβ42 toxicity.

Cytochrome c oxidase (COX), complex IV of the mitochondrial electron transport chain, uses energy from reducing molecular oxygen to water to generate a proton gradient across the inner mitochondrial membrane. Levels of mt:CoII (a COX subunit) declined in aged healthy control fly brain. mt:CoII expression was downregulated in Aβ42 flies compared to controls at all time-points and was stably-expressed across age following Aβ42 induction. The link between COX and AD is unclear, although Aβ is known to inhibit COX activity [38]. For example, in AD patients, COX activity—but not abundance—is reduced, resulting in increased levels of ROS [39]. However, in COX-deficient mouse models of AD, plaque deposition and oxidative damage are reduced [40]. Hence, the ageing-dependent decline in mt:CoII may represent either a reduction in COX function which renders the brain vulnerable to damage and is exacerbated by Aβ42 toxicity, or a protective mechanism against both ageing and amyloid toxicity.

The cuticle protein Acp65Aa was also upregulated in Aβ42 flies, but levels fell sharply between 5 and 19 days. However, it is surprising that we identified Acp65Aa in our samples, as it is not expected to be expressed in the brain. One explanation may involve chitin, which has been detected in AD brains and has been suggested to facilitate Aβ nucleation [41]. Amyloid aggregation has previously been shown to plateau around 15 days post-induction [42], which is around the same time that Acp65Aa drops in Aβ42 flies. Our results suggest that Aβ42 causes an increase in Acp65Aa expression early in the disease, but further experiments are needed to confirm this and to investigate its relationship with nucleation and the aggregation process.

#### Hub-bottlenecks: Hsp70A, Gp93, Top2 and Act75B

The four hub-bottlenecks are consistent with Aβ42 inducing stress. Hsp70A, a heat shock protein that responds to hypoxia, was significantly upregulated at early time-points (5 days) in Aβ42 flies, compared to healthy controls which exhibited stable expression of this protein throughout life. Although the levels dropped in Aβ42 flies between days 5 and 31 post-induction, at later time-points Hsp70A increased again, possibly suggesting a two-phase response to hypoxia in Aβ42 flies. We found that Gp93—a stress response protein that binds unfolded proteins—to be increased across age in Aβ42 flies compared to controls possibly suggesting an early and sustained protective mechanism against Aβ42-induced damage. DNA topoisomerase 2 (Top2), an essential enzyme for DNA double-strand break repair, was decreased in Aβ42 flies, following a pattern which mirrors changes in its expression with normal ageing. Double-strand breaks occur naturally in the brain as a consequence of neuronal activity—an effect that is aggravated by Aβ[7]. As a consequence of deficient DNA repair machinery, deleterious genetic lesions may accumulate in the brain and exacerbate neuronal loss.

Finally, we found that actin (Act57B) was increased in Aβ42 flies, in agreement with two recent studies on mice brains [43,44]. Kommaddi and colleagues found that Aβ causes depolymerisation of F-actin filaments in a mouse AD model before onset of AD pathology [44]. The authors showed that although the concentration of monomeric G-actin increases, the total concentration of actin remains unchanged. It has long been known that G-, but not F-, actin is susceptible to cleavage by trypsin [45], permitting its detection and quantification by IM-DIA-MS. Hence, the apparent increase of actin in Aβ42 flies may be due to F-actin depolymerisation, which increases the pool of trypsin-digestible G-actin, and is consistent with the findings of Kommaddi *et al*. To confirm whether total actin levels remain the same in the brains of Aβ42 flies, additional experiments would have to be carried out in the future, for example tryptic digestion in the presence of MgADP—which makes F-actin susceptible to cleavage [46]—and transcriptomic analysis of actin mRNA. Furthermore, actin polymerisation is ATP-dependent, so increased levels of G-actin may indicate reduced intracellular ATP. In addition, ATP is important for correct protein folding and therefore reduced levels may lead to increased protein aggregation in AD.

Due to the importance of these hub and bottleneck proteins in the protein interaction network, we predict that AD-associated alterations in their abundance will likely have a significant effect on the cellular dynamics of the brain.

### Dysregulated genes are associated with known AD and ageing network modules

Finally, we clustered the protein interaction network into modules and performed a Gene Ontology enrichment analysis on modules that contained any of the 228 significantly altered proteins. We saw no Gene Ontology term enrichment when we tested these proteins clustered according to their abundance profiles (Fig 2C), presumably because the proteins affected in AD are diverse and involved in many different biological processes. However, by testing network modules for functional enrichment, we exploited the principle that interacting proteins are functionally associated. Using a subgraph of the STRING network containing the significantly altered proteins and their directly-interacting neighbours, we used MCODE [47] to find modules of densely interconnected nodes. We chose to include neighbouring proteins to compensate for proteins that may not have been detected in the MS experiments due to the stochastic nature of observing peptides and the wide dynamic range of biological samples [48]. The resulting subgraph contained 4842 proteins, including 183 of the 228 significantly altered proteins, as well as 477 proteins that were only identified in healthy or Aβ42 flies and 3125 proteins that were not identified in our IM-DIA-MS experiments. 12 modules were present in the network (Fig 5A, Supplementary Data 2). The proportion of these modules that were composed of significantly altered proteins ranged from 0–8%. All but one of the modules were enriched for processes implicated in AD and ageing (Fig 5, Supplementary Data 3), including respiration and oxidative phosphorylation, transcription and translation, proteolysis, DNA replication and repair, and cell cycle regulation. These modules contained two proteins that were recently found to be significantly altered in the brain of AD mice [43] and are both upregulated four-fold in AD: adenylate kinase, an adenine nucleotide phosphotransferase, and the armadillo protein Arm, involved in creating long-term memories.

**Figure 5.**
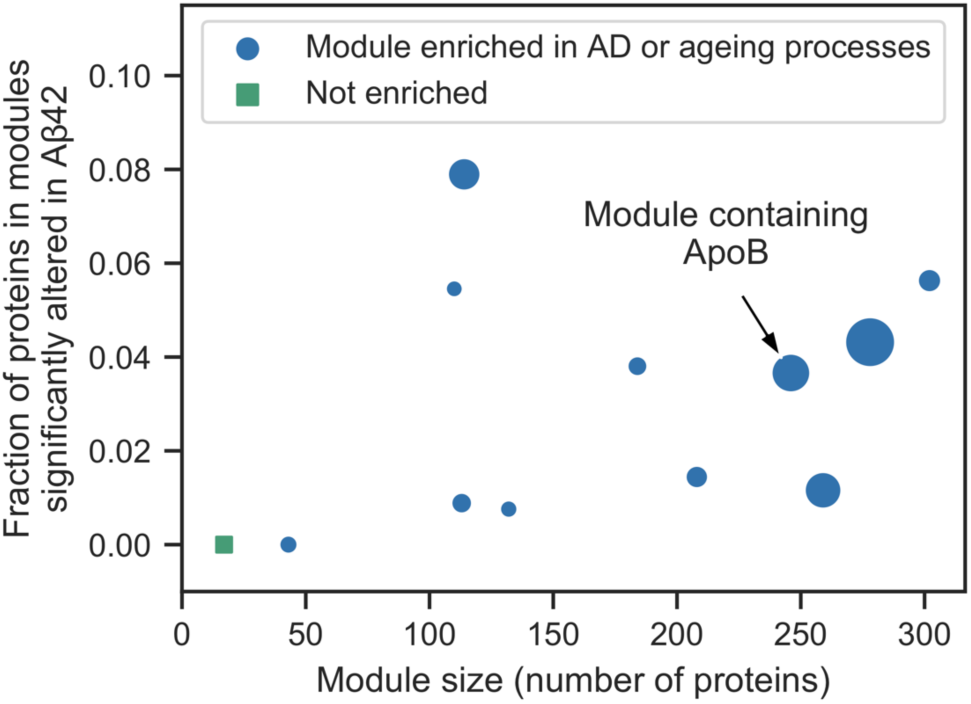
Analysis of network modules enriched for AD or ageing processes. MCODE was used to identify network modules in a subgraph of the STRING network containing the significantly altered proteins and their directly-interacting neighbours. The size of the resulting 12 modules is plotted against the fraction of proteins in these modules that are significantly altered in AD. Module 2 is annotated as containing ApoB. Marker sizes denote the MCODE score for the module.

In humans, the greatest genetic risk factor for AD is the Ɛ4 allele of ApoE—an apolipoprotein involved in cholesterol transport and repairing brain injuries [49]. A recent study showed that ApoE is only upregulated in regions of the mouse brain that have increased levels of Aβ [43], indicating a direct link between the two proteins. Although flies lack a homolog of ApoE, they do possess a homolog of the related apolipoprotein ApoB (Apolpp) [50], which contributes to AD in mice [51,52] and is correlated with AD in humans [53,54]. Interestingly, whilst it was not identified by IM-DIA-MS, ApoB interacts with 12 significantly altered proteins in the STRING network, so is included in the subgraph induced on the significantly altered proteins and their neighbours. ApoB was found in the second highest scoring module that contains proteins involved in translation and glucose transport (Fig 5) [55].

We analysed the 31 proteins significantly altered in normal ageing, but not AD. Of the 29 proteins that were contained in the STRING network, 24 interact directly with at least one of the AD significantly altered proteins, suggesting an interplay between ageing and AD at the pathway level. Using a subgraph of the STRING network induced on these proteins and their 1603 neighbours, we identified eight network modules that were enriched for ageing processes [56], including respiration, unfolded protein and oxidative damage stress responses, cell cycle regulation, DNA damage repair, and apoptosis.

## Discussion

Despite the substantial research effort spent on finding drugs against AD, effective treatments remain elusive. We need to better understand the molecular processes that govern the onset and progression of the complex pathologies observed in AD. This knowledge will help to identify new drug targets to treat and prevent AD.

Analysis of post-mortem human brain tissue is an important way to study dementia, but cannot capture the progression of pathology from the initiation of disease. Due to their short lifespan and ease of genetic manipulation, model organisms such as *Drosophila melanogaster* provide a tractable system in which to examine the progression of AD pathology across life. We performed a longitudinal study of the *Drosophila* brain proteome, using an inducible model of AD, label-free quantitative IM-DIA-MS and network analyses. We were able to track alterations in protein levels from the point of exposure to human Aβ42 and the widespread interaction of Aβ42 with brain signalling networks as pathology progresses.

Our proteomic analyses identified Aβ42-induced alterations in levels of 228 proteins, which clustered into four groups: those which were either elevated (cluster 1) or reduced (cluster 2) in AD relative to controls throughout life, those which were altered in correlation with ageing in healthy and Aβ42 flies (cluster 3), and those which changed in Aβ42 flies across life but independently of ageing-dependent effects in healthy controls (cluster 4). Further computational analysis of these proteins revealed significant network properties within the fly brain proteome. Assessing hub and bottleneck properties, many of the Aβ42-induced proteomic changes represented alterations in bottleneck proteins suggesting that they play key roles in downstream cellular function. Of these, some display non-hub properties indicating that they are important for maintaining cellular homeostasis in a targeted fashion, whereas others also displayed hub properties suggesting that they are central in linking cellular signalling pathways to maintain cell function.

We identified five nonhub-bottleneck proteins and four hub-bottleneck proteins, the expression of which was altered in Aβ42 flies relative to controls across life. Due to the importance of these hub and bottleneck proteins in the protein interaction network, we predict that AD-associated alterations in their abundance will likely have a significant effect on the cellular dynamics of the brain. Indeed, these proteins play key molecular roles in metabolism (AscI, Echs1, Got2), protein homeostasis (Hsp70A, Gp93), and protection against oxidative stress (mt:CoII) and DNA damage (Top2). These processes have been shown to affect neuronal function and protection against proteo-toxicity. Alterations in these proteins may represent either adaptive responses to the presence of abnormal protein aggregates, such as Aβ42, or mediators of neuronal toxicity. Further functional genomic studies are therefore required to establish the causal role of these processes in governing onset and progression of AD pathology.

Assessing the human orthologs of these genes, identified using DIOPT [57], indicates that several of these bottleneck proteins have been previously implicated in association with AD or other neurological conditions in humans or mammalian models of disease. ACSL4 (Acs1 ortholog) has been shown to associate with synaptic growth cone development and mental retardation [58]. Mutations in ECHS1 (Echs1 ortholog), an enzyme involved in mitochondrial fatty acid oxidation, associate with Leigh Syndrome, a severe developmental neurological disorder [59]. Proteomic studies have revealed that GOT2 (Got2 ortholog) is down-regulated in infarct regions following stroke [60], and in AD patient brain [61]. Integrating data from human post-mortem brain studies, HSPA1A (Hsp70Aa ortholog) upregulates in the protein interaction network of AD patients compared to healthy controls [62], and has recently been suggested to block APP processing and Aβ production in mouse brain [63]. Synthetic, fibrillar, Aβ42 reduces expression of TOP2B (Top2 ortholog) in rat cerebellar granule cells and in a human mesenchymal cell line, suggesting this may contribute to DNA damage in response to amyloid [64]. HSP90B1 (Gp93 ortholog) shows increased expression following TBI in mice [65], and associates with animal models of Huntington’s disease [66]. Finally, ACTB (Act57B ortholog) has been implicated as a significant AD risk gene and central hub node using integrated network analyses across GWAS [67].

ACSL4, ECHS1, and HSP90B1 have no reported association with AD or related dementias, however, which suggests that our study has potential to identify new targets in the molecular pathogenesis of this disease. Our study also provides additional information about the homeostasis of these proteins across life from the point of amyloid production. For example, the abundances of Acs1 and Got2 are elevated following Aβ42 induction and continue to increase with age relative to controls. Echs1 is reduced in Aβ42 flies compared to controls but increases across life in parallel with ageing-dependent increases in this protein. Structural proteins Acp65Aa and Act57B are elevated in response to Aβ42 but decline across life whilst remaining stable in control flies. Gp93 and Top2 are either elevated or reduced in response to Aβ42 but mirror ageing-dependent alterations in their expression. mt:CoII is reduced following Aβ42 expression at all time-points, but reduced with ageing in controls. Hsp70A is increased early in Aβ42 flies, reduced to control levels in mid-life then elevated at late pathological stages whilst remaining stable in healthy controls.

Analysing Gene Ontology enrichment using network modules, to capture the diverse biological processes modified in AD, we identified 12 modules enriched for processes previously implicated in ageing and AD. This validates the use of our *Drosophila* model in identifying progressive molecular changes in response to Aβ42 that are likely to correlate with progression of cognitive decline in human disease. Further work is required to modify the genes identified in our study at different ages, in order to elucidate whether they represent mediators of toxicity as disease progresses, factors which increase neuronal susceptibility to disease with age or compensatory protective mechanisms. Model organisms will be essential in unravelling these complex interactions. Our study therefore forms a basis for future analyses that may identify new targets for disease intervention that are specific to age and/or pathological stage of AD.

## Materials and methods

### Fly stocks

The TgAD fly line used in this study [23] contains the human transgene encoding the Arctic mutant Aβ42 peptide under the control of an Upstream Activation Sequence (UAS) [68]. Expression of Aβ42 was controlled by GeneSwitch [69]—a mifepristone-inducible GAL4/UAS expression system—under the pan-neuronal elav promoter. All flies were backcrossed for six generations into the w^1118^ genetic background.

Flies were grown in 200 ml bottles on a 12 h/12 h light/dark cycle at constant temperature (25 °C) and humidity. Growth media contained 15 g/l agar, 50 g/l sugar, 100 g/l autolysed yeast, 100 g/l nipagin and 3 ml/l propionic acid. Flies were maintained for two days after eclosion before females were transferred to vials at a density of 25 flies per vial for the lifespan analysis and 10 flies per vial for the IM-DIA-MS analysis. Expression of Aβ42 was induced in TgAD flies by spiking the growth media with mifepristone to a final concentration of 200 µM. Flies were transferred to fresh media three times per week, at which point the number of surviving flies was recorded. For each of the three biological repeats, 10 healthy and 10 Aβ42 flies were collected at 5, 19, 31 and 46 days, as well as 54 and 80 days for healthy flies. Following anesthetisation with CO_2_, brains were dissected in ice cold 10 mM phosphate buffered saline snap frozen and stored at −80°C.

### Extraction of brain proteins

Brain proteins were extracted by homogenisation on ice into 50 µl of 50 mM ammonium bicarbonate, 10 mM DTT and 0.25% RapiGest detergent. Proteins were solubilised and disulfide bonds were reduced by heating at 80°C for 20 minutes. Free cysteine thiols were alkylated by adding 20 mM IAA and incubating at room temperature for 20 minutes in darkness. Protein concentration was determined and samples were diluted to a final concentration of 0.1% RapiGest using 50 mM ammonium bicarbonate. Proteins were digested with trypsin overnight at 37°C at a 50:1 protein:trypsin ratio. Additional trypsin was added at a 100:1 ratio the following morning and incubated for a further hour. Detergent was removed by incubating at 60°C for 1 hour in 0.1% formic acid. Insoluble debris was removed by centrifugation at 14,000 × g for 30 minutes. Supernatant was collected, lyophilised and stored at −80°C. Prior to lyophilisation peptide concentration was estimated by nanodrop (Thermo Fisher Scientific, Waltham, MA).

### Label-free quantitative IM-DIA-MS

Peptides were separated by nanoscale liquid chromatography (LC) by loading 300 ng of protein onto an analytical reversed phase column. IM-DIA-MS analysis was performed using a Synapt G2-Si mass spectrometer (Waters Corporation, Manchester, UK). The time-of-flight analyzer of the instrument was externally calibrated with a NaCsI mixture from m/z 50 to 1990. Spectra were acquired over a range of 50–2000 m/z. Each biological repeat was analysed at least twice to account for technical variation.

LC-MS data were peak detected and aligned by Progenesis QI for proteomics (Waters Corporation). The principles of the embedded search algorithm for DIA data has been described previously [70]. Proteins were identified by searching against the *Drosophila melanogaster* proteome in UniProt, appended with common contaminants, and revered sequence entries to estimate protein identification false discovery rate (FDR) values, using previously specified search criteria [71]. Peptide intensities were normalised to control for variation in protein loading and relative quantification. Abundances were estimated by Hi3-based quantitation [72].

### Data analysis

Proteins that were identified in both healthy and Aβ42 flies were considered for further analysis. Missing data were replaced by the minimum abundance measured for any protein in the same repeat [48]. The data were quantile normalised [73], so that different conditions and time points could be compared reliably. Quantile normalisation transforms the abundances so that each repeat has the same distribution.

For PCA analysis, the data were log_10_-transformed and each protein was standardised to zero mean and unit variance. Hierarchical biclustering was performed using the Euclidean distance metric with the complete linkage method. Prior to clustering, proteins were normalised to their abundance in healthy flies at 5 days.

Proteins that were identified by IM-DIA-MS in either healthy or Aβ42 flies were assessed for overrepresentation of Gene Ontology terms using GOrilla [74], which uses ranked lists of target and background genes. Proteins were ranked in descending order by their mean abundance. The type I error rate was controlled by correcting for multiple testing using the Benjamini-Hochberg method at an FDR of 5%.

Clusters of proteins were assessed for overrepresentation of GO-Slim terms in the Biological Process ontology using Panther (version 13.1) with a custom background of the 3093 proteins identified by IM-DIA-MS in healthy or AD flies.

### Identification of significantly altered proteins

Significantly altered proteins were identified using five methods that are frequently used to identify differentially expressed genes in time course RNA-Seq data. DESeq2 [31], EDGE [33], edgeR [30], limma [32] and maSigPro [34] are all available in R through Bioconductor. Dispersions were estimated from the biological and technical repeats. Unless otherwise stated, default parameters were used for all methods under the null hypothesis that a protein does not change in abundance between healthy and AD conditions in normal ageing. The type I error rate was controlled by correcting for multiple testing using the Benjamini-Hochberg method at a FDR of 5%. A protein was classified as significantly altered if two or more methods identified it.

DESeq2 models proteins with the negative binomial distribution and performs likelihood ratio tests. A time course experiment was selected in EDGE using the likelihood ratio test and a normal null distribution. edgeR uses the negative binomial distribution and performs quasi-likelihood tests. limma fits linear models to the proteins and performed empirical Bayes F-tests. maSigPro fits generalised linear models to the proteins and performs log-likelihood ratio tests.

Significantly altered proteins were clustered using a Gaussian mixture model. Protein abundances were log10-transformed and *z* scores were calculated. Gaussian mixture models were implemented for 1–228 clusters. The best model was chosen using the Bayesian information criterion (BIC), which penalises complex models:

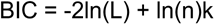

where ln(L) is the log-likelihood of the model, n is the number of significantly altered proteins and k is the number of clusters. The model with lowest BIC was chosen.

### Networks

All network analysis was performed using the *Drosophila melanogaster* STRING network (version 10) [35]. Low confidence interactions with a ‘combined score’ < 500 were removed in all network analyses.

Network properties of the significantly altered proteins were analysed in the brain protein interaction network. A subgraph of the STRING network was induced on the 3093 proteins identified by IM-DIA-MS in healthy or Aβ42 flies and the largest connected component was selected (2428 nodes and 44,561 edges). The subgraph contained 183 of the 228 significantly altered proteins. For these proteins, four network properties were calculated as test statistics: mean node degree; mean unweighted shortest path length between a node and the remaining 182 nodes; the size of the largest connected component in the subgraph induced on these nodes; and mean betweenness centrality. Hypothesis testing was performed using the null hypothesis that there is no difference between the nodes in the subgraph. Assuming the null hypothesis is true, null distributions of each test statistic were simulated by randomly sampling 183 nodes from the network 10,000 times. Using the null distributions, non-parametric one-sided p-values were calculated as the probability of observing a test statistic as extreme as the test statistic for the significantly altered proteins.

A subgraph of the STRING network was induced on the proteins significantly altered in AD and their neighbours and the largest connected component was selected (4842 nodes and 182,474 edges). The subgraph contained 198 of the 228 significantly altered proteins and was assessed for enrichment of Gene Ontology terms. Densely connected subgraphs were identified using MCODE [47]. Modules were selected with an MCODE score > 10. As STRING is a functional interaction network, clusters of nodes may correspond to proteins from the same complex, pathway or functional family. Clusters were assessed for overrepresentation of GO-Slim terms in the Biological Process ontology using Panther (version 13.1) [75] with a custom background of the 3093 proteins identified by IM-DIA-MS in healthy or Aβ42 flies. Fisher’s exact tests were performed and the type I error rate was controlled by correcting for multiple testing using the Benjamini-Hochberg method at a FDR of 5%.

### Open source software

Data analysis was performed in Python 3.6 (Python Software Foundation, http://www.python.org) using SciPy [76], NumPy [77], Pandas [78], scikit-learn [79], NetworkX [80], IPython [81] and Jupyter [82]. Figures were plotted using Matplotlib [83] and seaborn.

## Acknowledgements

We thank Dr Damian Crowther (University of Cambridge) for donation of UAS-Aβ42 fly stocks and Dr Hervé Tricoire (CNRS, France) for donation of elavGS fly stocks. Fig 1C: fly graphic by Daan Kauwenberg and brain graphic by Julia Amadeo, both from the Noun Project.

## Funding

H.S. is supported by an ISMB Wellcome PhD studentship [203780/Z/16/A]; A.C. is supported by a BBSRC CASE PhD studentship and Waters Corporation; J.L. is supported by BBSRC [BB/L002817/1]; L.P. and F.K. are supported by a Wellcome Trust Strategic Award to L.P. [098565/Z/12/Z] and an Alzheimer’s Research UK Project Grant (ART-2009-4). The work was supported by a Wellcome instrumentation grant [104913/Z/14/Z].

## Competing interests

None

## Supplementary Information

### Methods

#### IM-DIA-MS analysis

Nanoscale liquid chromatography (LC) separation of tryptic peptides was performed using a nanoAcquity UPLC system (Waters Corporation) equipped with a UPLC HSS T3 1.7 µm, 75 µm × 250 mm analytical reverse phase column (Waters Corporation). Prior to peptide separation, 300 ng of tryptic peptides were loaded onto a 2G, V/V 5 µm, 180 µm × 20 mm reverse phase trapping column at 5 µl/min for 3 minutes. IM-DIA-MS analysis of tryptic digests was performed using a Synapt GS-Si mass spectrometer equipped with a T-Wave-IMS device. Mass measurements were made in positive-mode ESI with the instrument operated in resolution mode with a typical resolving power of 20,000 full width at half maximum. Prior to analysis the time-of-flight analyzer was externally calibrated with a NaCsI mixture from *m/z* 50 to 1990. The data were post-acquisition lock mass corrected using the double charged monoisotopic ion of [Glu1]-Fibrinopeptide B. To achieve lock mass correction, a 100 fmol/µl solution of [Glu1]-Fibrinopeptide B was infused at a 90° angle to the analytical sprayer. This reference sprayer was sampled every 60 seconds. Accurate IM-DIA-MS data were collected in the DIA mode of analysis, HDMS^E^ [24,71] IM spectrometry was performed by applying a constant wave height of 40 V whilst a constant wave velocity of 650 m/s was maintained. Wave heights within the trap and transfer were both set at 4 V whilst the wave velocities were 311 and 175 m/s respectively. MS data were acquired over 50-2000 *m/z for* each mode. Spectral acquisition time for each mode was 0.5 s with a 0.015 interscan delay, corresponding to a cycle of low and elevated energy data being acquired every 1.1 s. During the low energy MS mode data was acquired whilst applying a constant collision energy of 4 eV within the transfer. After IMS, MS/MS data was acquired by ramping the collision energy within the transfer region between 15 and 45 eV. To ensure that ions with a *m/z* less than 350 were derived from peptide fragmentation within the transfer region the radio frequency applied to the quadrupole mass analyser was adjusted to optimise transmission within the region of 350 – 2000 Da. Each biological replicate was analysed at least twice.

#### MS Data Processing

All MS data were processed in Progenesis QI for proteomics. Data were imported into Progenesis to generate a 3D representation of the data (*m/z*, RT and peak intensity). Samples were then time aligned with the software allowed to automatically determine the best reference run from the dataset. Following alignment, peak picking was performed on MS level data. A peak picking sensitivity of 4 (out of 5) was set. Peptide features were tentatively aligned with their respective fragment ions based primarily on the similarity of their chromatographic and mobility profiles. Requirements for features to be included in post-processing database searching were as follows: 300 counts for low energy ions, 50 counts for high energy ions and 750 counts for deconvoluted precursor intensities. Subsequent data were searched against 20,049 sequences from the UniProt canonical *Drosophila* database (appended with common contaminants). Trypsin was specified as the enzyme of choice and a maximum of two missed cleavages were permitted. Carbamidomethyl (C) was set as a fixed modification whilst oxidation (M) and N-terminal acetylation were set as variable modifications. Peptide identifications were grouped and relative quantification was performed using non-conflicting peptides only.

### Data

**Supplementary Data 1**

supplementary_data_1.xlsx

Proteomics data

**Supplementary Data 2**

supplementary_data_2.txt

MCODE modules

**Supplementary Data 3**

supplementary_data_3.xlsx

Gene Ontology enrichment

**Figure S1:**
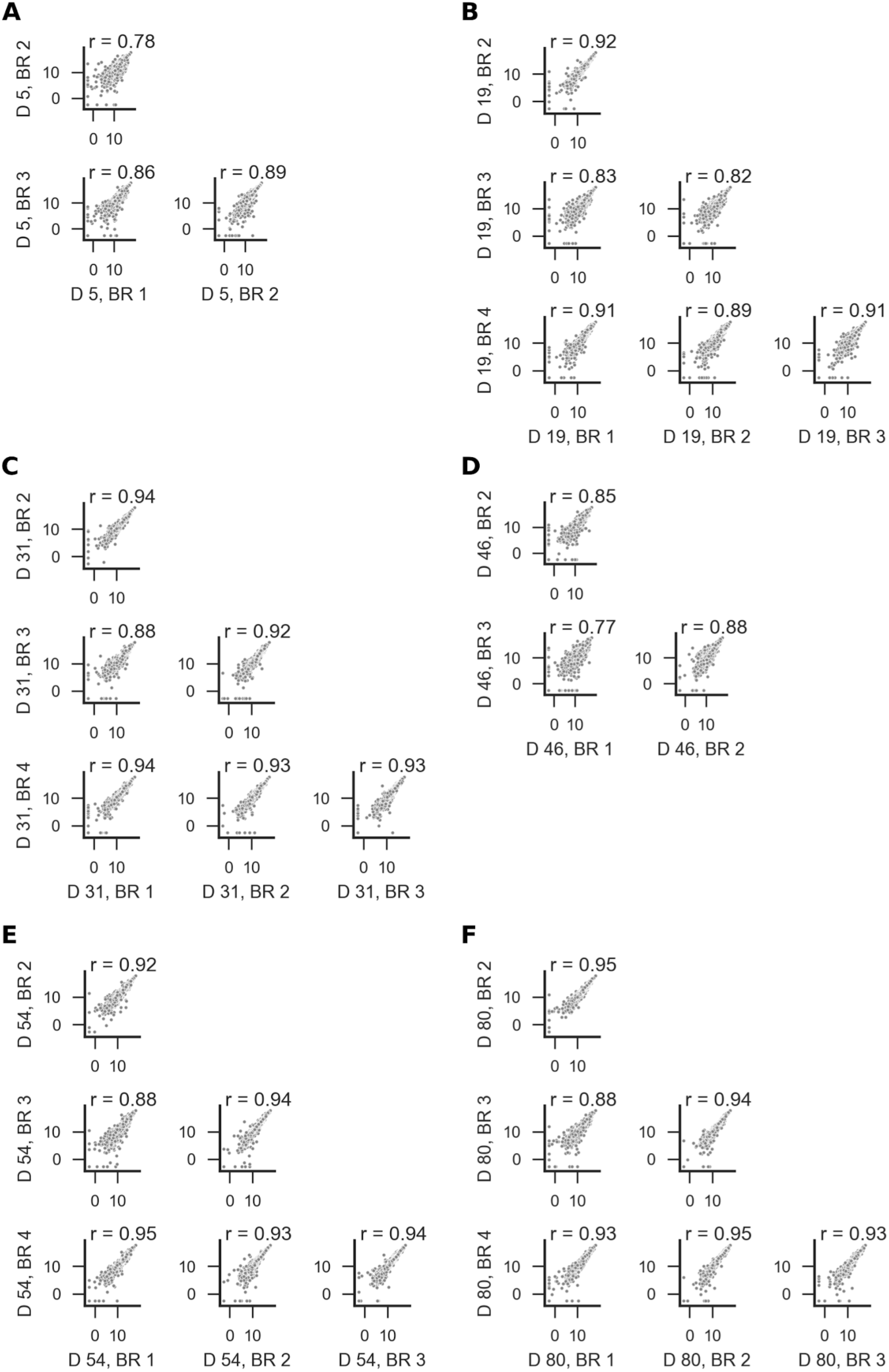
Assessment of experimental reproducibility. Scatter plots comparing protein abundances in different biological repeats (BR) of healthy flies at days (D) (**A**) 5, (**B**) 19, (**C**) 31, (**D**) 46, (**E**) 54 and (**F**) 80. Abundances were log2-transformed before plotting. Pearson correlation coefficients (r) are shown for each pair of biological repeat at each time point.

**Figure S2:**
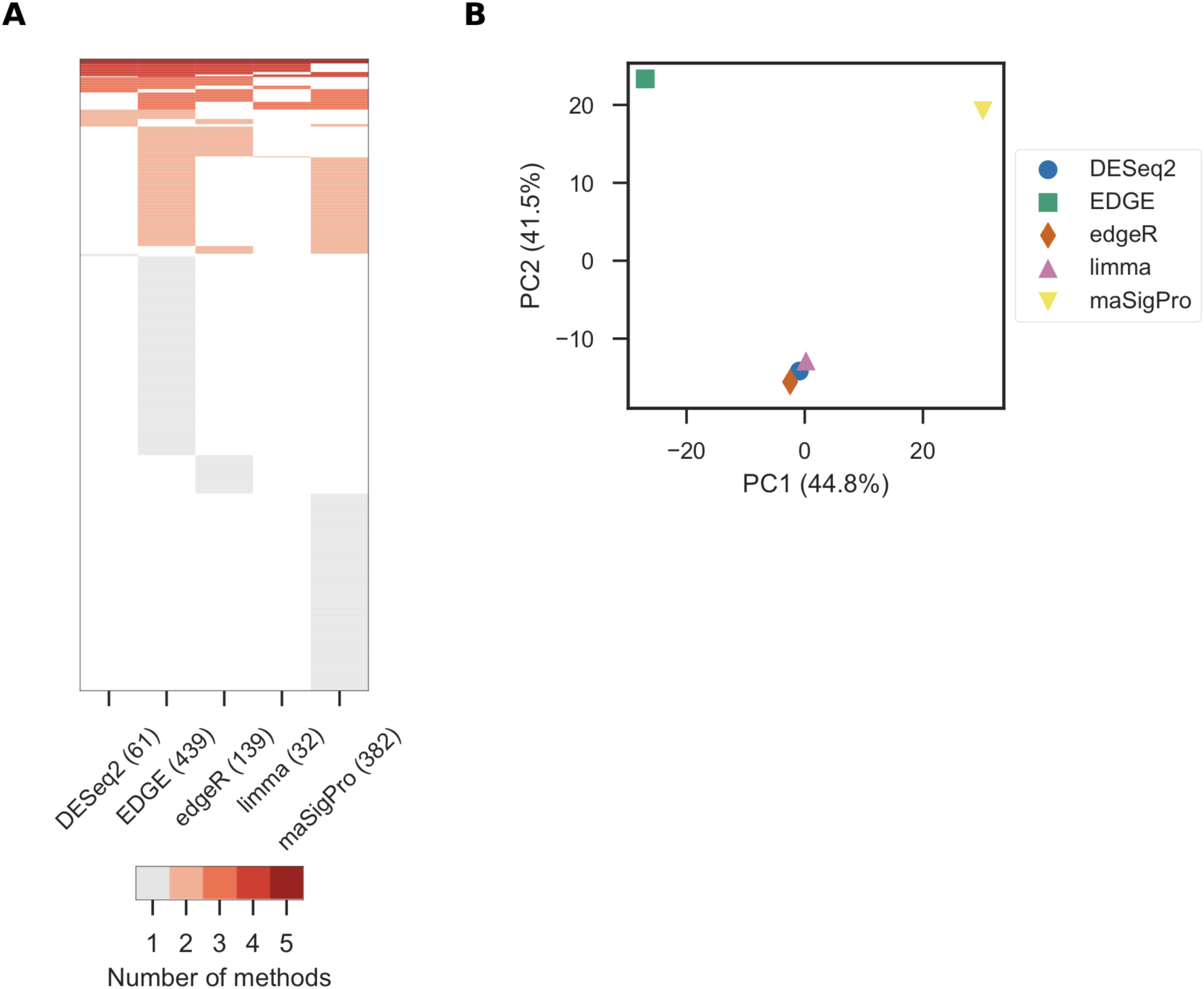
Analysis of the five statistical methods used to identify significantly altered proteins. (**A**) Heat map of the proteins detected by each method. (**B**) Principal component analysis of these results. Axes are annotated with the percentage of variance explained by each principal component.

**Figure S3:**
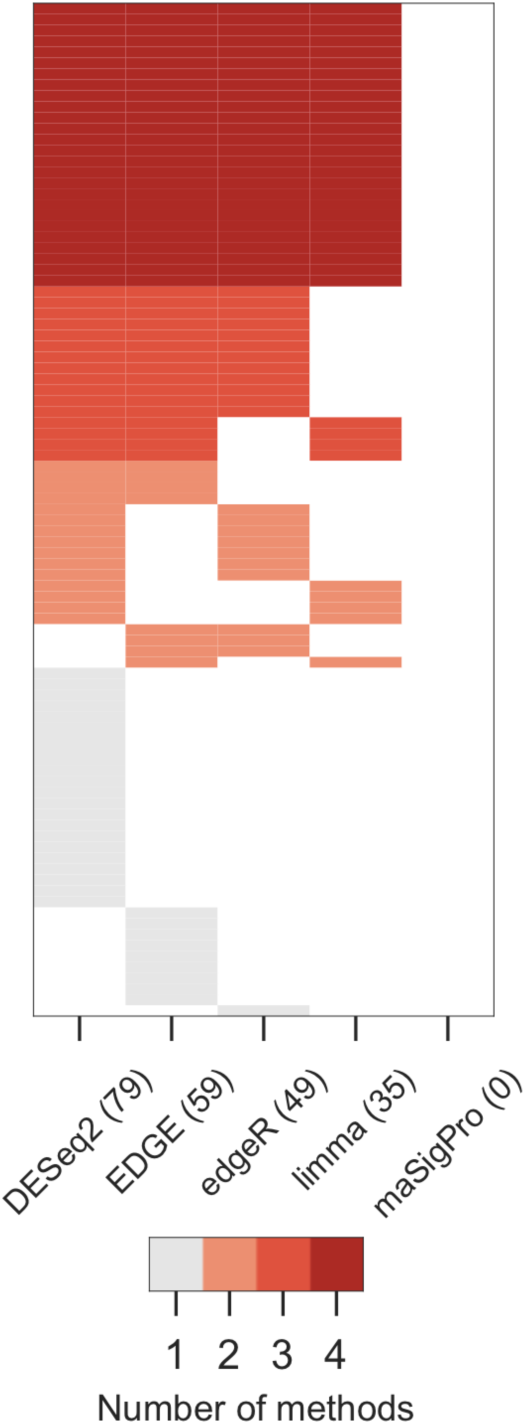
Identification of significantly altered proteins during normal ageing. Heat map of the proteins detected by each method.

**Figure S4:**
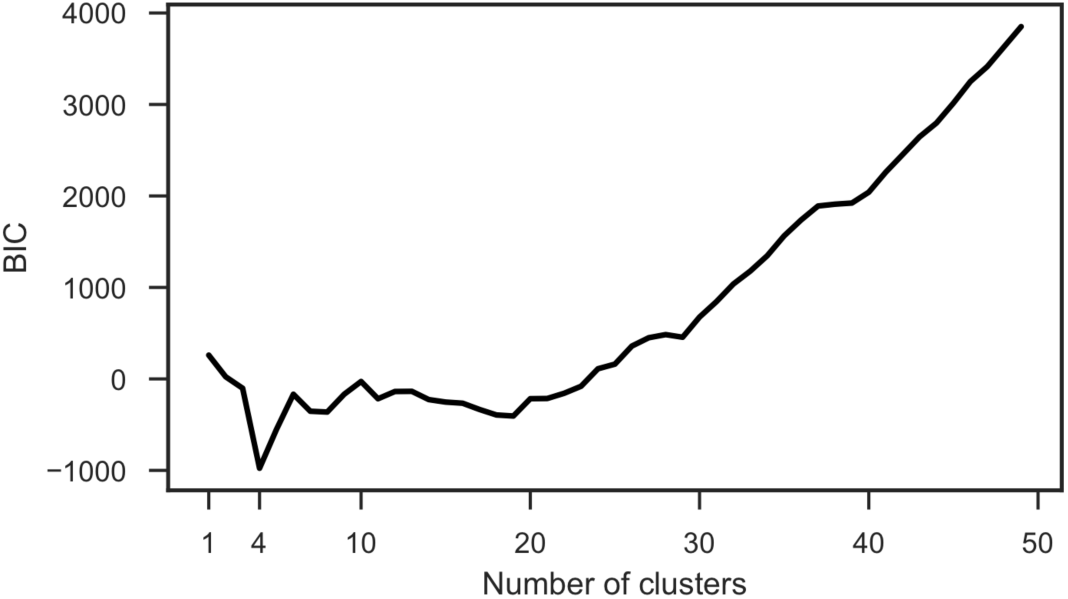
Model selection for clustering of the significantly altered proteins using a Gaussian mixture model. The best model was chosen using the Bayesian information criterion (BIC), which penalises complex models.

